# An integrative process-based model for biomass and yield estimation of hardneck garlic (*Allium sativum*)

**DOI:** 10.1101/2021.11.04.467009

**Authors:** Kyungdahm Yun, Minji Shin, Kyung Hwan Moon, Soo-Hyung Kim

**Affiliations:** University of Washington; Research Institute of Climate Change and Agriculture

## Abstract

We introduce an integrative process-driven crop model for garlic (*Allium sativum*). Building on our previous model that simulated key phenological, morphological, and physiological features of a garlic plant, the new garlic model provides comprehensive and integrative estimations of biomass accumulation and yield formation under diverse environmental conditions. This model also showcases an application of Cropbox to develop a comprehensive crop model. Cropbox is a crop modeling framework featuring declarative modeling language and unified simulation interface for building and improving crop models. Using Cropbox, we first evaluated the model performance against three datasets with an emphasis on biomass and yield measured under different environmental conditions and growing seasons. We then applied the model to simulate optimal planting dates under future climate conditions for assessing climate adaptation strategies between two contrasting locations in South Korea: the current growing region (Gosan) and an unfavorable cold winter region (Chuncheon, Gangwon-do). The model simulated the growth and development of a southern-type cultivar (Namdo, Jeju-do) reasonably well. Under RCP (Representative Concentration Pathway) scenarios, an overall delay in optimal planting date from a week to a month and a slight increase in potential yield were expected in Gosan. Expansion of growing region to northern area including Chuncheon was expected due to mild winter temperatures in the future and may allow Namdo cultivar production in more regions. The predicted optimal planting date in the new region was similar to the current growing region that favors early fall planting. Our new integrative garlic model provides mechanistic, process-driven crop responses to environmental cues and can be useful for assessing climate impacts and identifying crop specific climate adaptation strategies for the future.

## 1 Introduction

Garlic (*Allium sativum*) is a historically important horticultural crop in many countries with global production reaching 30.7 million tons in 2019 after 40% increase of production in the last decade (FAO, 2020). Physiology of garlic has been extensively studied with an emphasis on characteristics as a bulbous crop where a clove of bulb is planted for growth and a newly grown bulb is harvested for storage and the next round of planting (Takagi, 1989; Kamenetsky, 2007). Some knowledge has been transferred to building crop models specifically targeted for simulating garlic growth and estimating yield at harvest. An early attempt for building a whole-plant garlic model was based on radiation-use efficiency (RUE) to obtain the total amount of carbon assimilates (Rizzalli et al., 2002). CropSyst, which is also a crop model based on RUE, was parameterized for garlic and used for simulating crop rotation between garlic and wheat (Giménez et al., 2016). Other studies focused on a certain aspect of garlic growth and development. The crop coefficient (*K_c_*) needed for calculating evapotranspiration with Penman-Monteith equation was specifically determined for garlic (Villalobos et al., 2004). Photosynthesis and transpiration responses to various environmental conditions were obtained for building a leaf-level gas-exchange model for garlic (Kim et al., 2013). Photosynthetic responses under elevated CO_2_ and nitrogen fertilization were further investigated for building a robust model for future climate conditions (Nackley et al., 2016). With a coupled gas-exchange model parameterized for garlic, a process-based model for simulating leaf development and growth of hardneck garlic was developed (Hsiao et al., 2019). Phenology of leaf initiation and appearance was individually tracked by taking account of multiple cues including thermal time accumulation, bulb storage effect, and photoperiod. Individual leaf elongation was translated and aggregated into leaf area expansion at canopy level which was then divided into two layers of sunlit and shaded leaves for accounting assimilated carbon based on the coupled gas-exchange model. However, carbon partitioning into plant organs such as bulb was not validated and thus not used for yield estimation. Carbon partitioning is a crucial step in yield estimation modeling for horticultural crops in a sense that the final yield is a result of biomass partitioned into a certain organ, such as bulb, to be harvested (Marcelis et al., 1998). The early garlic model used a set of multiple partitioning coefficients dynamically varying with developmental stages of the plant (Rizzalli et al., 2002).

Yield estimation for garlic cultivars grown in each region has been crucial to manage growing practices and maintain a stable supply in the market (Põldma et al., 2005; Lee et al., 2011; Abdalla et al., 2011; Portela et al., 2012). Regression models based on weather data or satellite images were often used for yield estimation at a large scale with a minimum set of historical input data (Choi and Baek, 2016; Gómez et al., 2021). Although worldwide garlic production has been steadily increased in the recent years, it is unclear if yield will be stable in the current growing regions under future climate conditions. For estimating yield under previously unobserved conditions, it is critical to develop a physiological understanding of how garlic would respond to these environmental cues (Nackley et al., 2016). Regional climate changes may lead to changes in current farming practices and shifts in growing regions to maintain or maximize the yield. Crop model is an important tool for supporting such decisions by enabling simulations of plant growth under diverse climate adaptation strategies (Rosenzweig et al., 2014; Holzkämper et al., 2015; Corbeels et al., 2018).

In this study, our primary objective was to build an integrative process-driven garlic model suitable for estimating harvestable biomass as scape and bulb yield under diverse environments including future climate conditions. A new model was developed based on an existing process-based garlic model with an original emphasis on phenology and extended with a focus on biomass accumulation and partitioning (Hsiao et al., 2019). The model was also improved to better reflect physiological responses to temperature by taking account storage conditions of seed garlic bulbs and cold stress response in terms of leaf-level growth and canopy-level mortality. We used Cropbox modeling framework for reimplementation to take advantage of its declarative modeling language and unified interface for coordinating a large batch of simulations with minimum configurations (Yun et al., 2020). For model testing, a new parameter set was specifically calibrated for Namdo cultivar (*Allium sativum* ‘Namdo’) and validated with multiple datasets with biomass measurements. For demonstrating yield estimation capability of the model, a climate adaptation strategy was assessed by model simulations for the same cultivar grown in two locations of South Korea. Namdo is a cultivar originally adapted to warm climate in southern region of Korea (Kim et al., 2009). However, the boundary of growth region between northern and southern types of garlic has moved northward in the past decades due to warming climate (Heo et al., 2006). Optimal planting dates for achieving maximum yield were discovered through model simulation under current and future climate conditions in the two locations where one is an already established region for growing southern-type garlic and the other has a potential to become a new establishment in the future.

## 2 Materials and Methods

### 2.1 Garlic Model

The garlic growth model was extended from a process-based model for leaf development and growth in hardneck garlic (Hsiao et al., 2019). The original model was capable of simulating leaf area expansion at an individual leaf level and estimating carbon assimilated in a canopy calculated by coupled gas-exchange, but assessing biomass allocated into a particular organ, *i.e.* bulb, was not a primary subject of the model at the time. For realistic yield prediction under future scenarios, the model should be able to be reliable under diverse environmental conditions (IPCC, 2014).

Model changes made for the experiments reported in this paper include improved biomass allocation, a dynamically adjusted phyllochron, and cold stress response (Figure 1). For the sake of technical convenience, model code originally written in C++ was reimplemented with Cropbox modeling framework using Julia programming language (Cropbox.jl, 2021). The Cropbox framework allowed a streamlined model development from model description in a concise declarative form, iterative parameter adjustments within a notebook environment, and batch simulation of large sets of parameters and production of figures included in this paper. The model source code is also available in the public repository (Garlic.jl, 2021).

**Figure 1:**
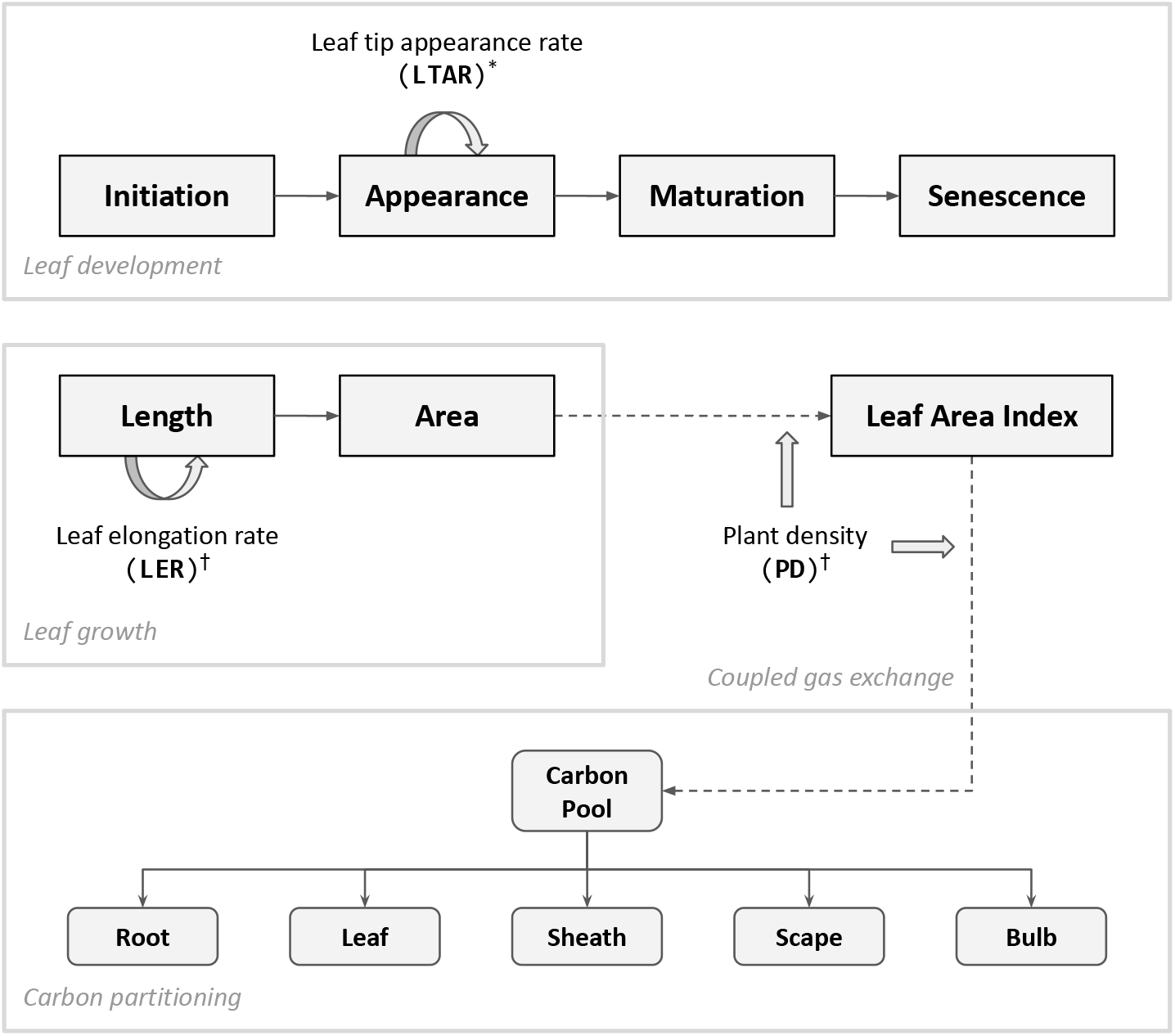
Diagram of key processes updated in the model presented in this paper. Leaf appearance is driven by accumulation of leaf tip appearance rate (LTAR) which is modified according to dynamic phyllochron (*). Cold stress (^†^) is implemented in two folds; individual leaf-level growth driven by leaf elongation rate (LER) and plant population-level response accounted by plant density (PD). Carbon assimilated through coupled gas exchange forms a carbon pool which determines the amount of carbon partitioned into individual organs. The bulb carbon constitutes a final yield of garlic. For a complete structure of the model, refer to Figure 1 of Hsiao et al. (2019).

#### 2.1.1 Biomass Allocation

The total amount of carbon assimilation was calculated by a C_3_ photosynthesis model coupled with a stomatal conductance model and energy balance model as described in the previous paper (Hsiao et al., 2019). Note that overall model structure including the gas-exchange module was reorganized for taking advantage of domain-specific language provided by Cropbox modeling framework (Yun et al., 2020). The assimilated carbon accumulates and forms a carbon pool (g) ready to be distributed to plant organs. A potential allocation rate of available carbon (gd^−1^) is driven by carbon supply rate from the pool (gd^−1^) excluding maintenance respiration (gd^−1^). After taking account of carbohydrate synthesis efficiency (*Y_g_*), an actual allocation rate (gd^−1^) is determined and split into a set of allocation rates for structural organs according to a partitioning table (Figure 2).

**Figure 2:**
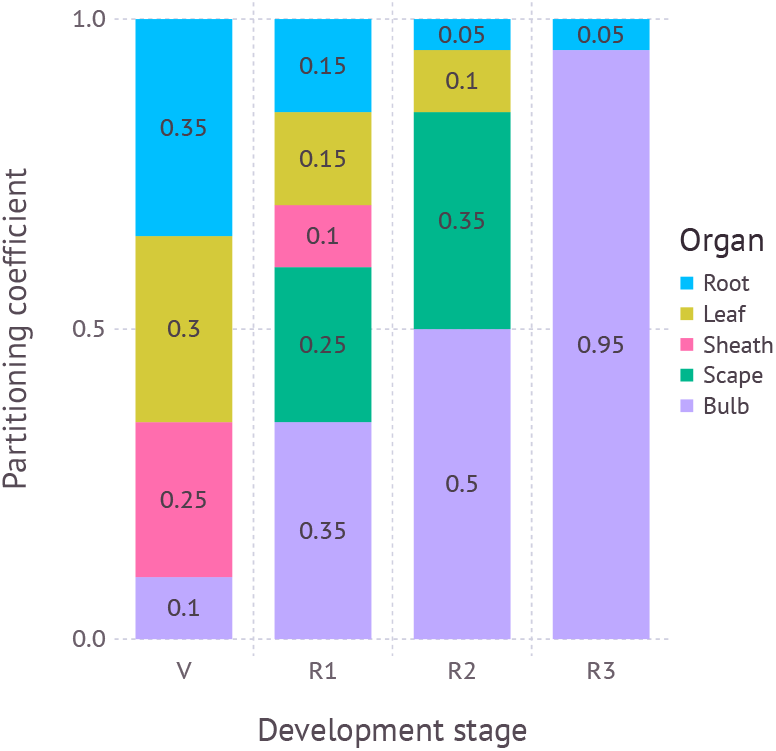
Partitioning coefficients for plant organs dynamically adjusted according to developmental stage. V: vegetative stage; R1: reproductive stage between scape initiation and appearance; R2: reproductive stage between scape appearance and removal; R3: reproductive stage after scape removal. R3 is not used if scape is not removed during simulation.

The partitioning table is a 2-dimensional array where each row represents a developmental stage and column represents a destination. Developmental stages span from seed, vegetative, bulb growth before scape appearance, bulb growth after scape appearance, bulb growth after scape removal, and death. Partitioning destinations include root, leaf, sheath, scape, and bulb. For each time step of simulation, an actual allocation rate weighted by a partitioning coefficient found in the table was used for biomass accumulation of each destination organ.

#### 2.1.2 Dynamic Phyllochron

An interval between leaf appearance, phyllochron, is not necessarily static throughout plant growth, but can dynamically change depending on the growth condition. However, maximum leaf tip appearance rate (LTAR_max_; d^−1^) in the original garlic model was determined by thermal time based on storage duration between harvest and planting as well as storage temperature for which seed garlic has been kept during this period. Once initialized, LTAR_max_ would have remained the same until the end of simulation. This assumption was often held when storage duration was close to an average duration where curve fitting was originally done for. However, sometimes the leaf tip appearance rate may have stayed too high at the end of growing season when storage duration was longer than usual. The opposite would happen when the storage duration was too short. In other words, scape appearance in the reproductive stage, which is driven by the same mechanism relying on phyllochron that assumes a scape appears after three phyllochrons since the onset of reproductive stage, could become too sensitive to the initial value of LTAR_max_. To alleviate this issue, LTAR_max_ was adjusted to converge towards a half of the asymptote of maximum leaf tip appearance rate parameter (LTAR_max,a_; d^−1^). The initial value LTAR_max,s_ is the same as maximal rate of leaf tip appearance rate modified by storage duration SD defined in the original model Hsiao et al. (2019). *SD_m_* is the storage duration that results into maximal leaf tip appearance rate and *α* controls steepness of the sigmoidal storage function. The same value of parameters were adopted from the original model (Table 1). After each leaf appearance, LTAR_max_ would linearly increase or decrease depending on the initial rate and the slope towards the asymptote. Convergence is done when leaf rank *k* reaches the generic leaf number *N_g_* which was set to 10 by default.

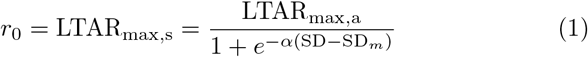

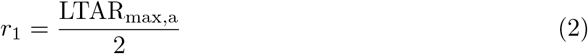

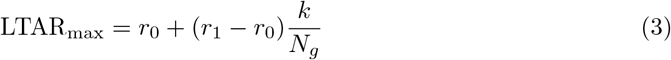

**Table 1:** Parameters for dynamic phyllochron

| Symbol | Value | Units | Description |
| --- | --- | --- | --- |
| $\text{LTAR}_{\text{max},a}$ | 0.4421 | $\text{d}^{-1}$ | Maximal leaf tip appearance asymptote |
| ST | 8 | $^{\circ}\text{C}$ | Storage temperature |
| $\text{SD}_m$ | 117.7523 | d | Storage days lead to maximal leaf tip appearance rate |
| $\alpha$ | 0.0256 | $\text{d}^{-1}$ | Steepness of sigmoidal storage function |
| $N_g$ | 10 | - | Generic leaf number |

#### 2.1.3 Cold Stress

Two types of cold stress response commonly found with garlic plants grown in the field during winter season were added to the model: cold injury and cold damage. Cold injury represents impeded leaf growth under below normal air temperatures. Actual leaf elongation rate (LER; cmd^−1^) is scaled down from potential leaf elongation rate (LER_p_; d^−1^) by cold injury effect (*E*) which is a function of the number of cold days (*D*) the plant has experienced. The number of cold days (D) increases when the air temperature (*T*) is below a critical temperature for cold injury (T_c,i_; °C). Once the temperature rises above the threshold T_c,i_, the cumulative days *D* decreases until it resets to zero. It assumes that the effect of cold injury is prolonged for a certain period time while recovering under normal temperature. The longer a plant is exposed to the cold temperature, the more difficult to recover from the injury. We assumed critical temperature where cold injury begins to appear was at 0 °C and had related factors (*a,b*) fitted with a dataset obtained from a growth chamber experiment (Table 2).

**Table 2:**
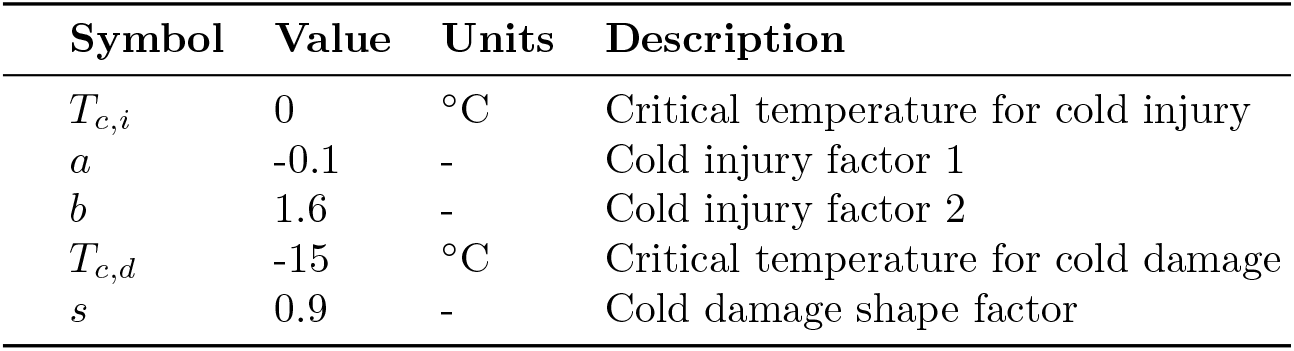
Parameters for cold stress response

| Symbol | Value | Units | Description |
| --- | --- | --- | --- |
| $T_{c,i}$ | 0 | °C | Critical temperature for cold injury |
| $a$ | -0.1 | - | Cold injury factor 1 |
| $b$ | 1.6 | - | Cold injury factor 2 |
| $T_{c,d}$ | -15 | °C | Critical temperature for cold damage |
| $s$ | 0.9 | - | Cold damage shape factor |

Namdo (ND) cultivar were planted and grown in pots under room temperature until the third leaf emerged. Plants were then placed in a growth chamber subject to constant temperature of 0 °C, −5 °C, −10 °C and −15 °C, respectively. Each treatment had three replicates and leaf area of the third leaf blade from survived plants was measured for eight days. Then cold injury effect (*E*) was fitted to the relative portion of survived plants after a given time.

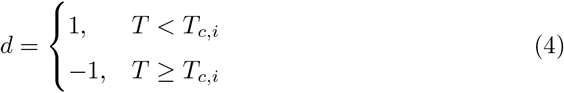

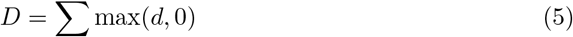

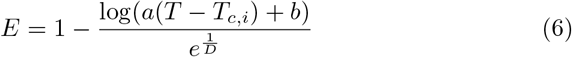

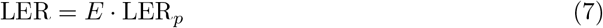

Cold damage represents death of the plant under more extreme temperature, resulting into a reduced plant density (PD). The current plant density (PD) starts from an initial planting density (PD_0_) and then may decrease over time as the air temperature (*T*) drops below a critical temperature for cold damage (*T_c,d_*). Unlike cold injury where leaf growth only slows down under low temperature and eventually recovers once temperature rises back up, cold damage leads to permanent wilting from which plants no longer recover. Mortality due to cold damage (*M*) is a logistic curve representing a relative portion of plants survived at the end of the cold damage treatments. Cold damage shape factor (*s*) determines how quickly the plants will die off under such extreme conditions. Due to lack of experimental datasets available for parameter fitting, we instead looked up episodes of cold damage and corresponding temperatures reported in newspaper during the past years to determine parameter values.

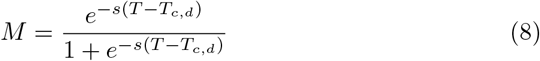

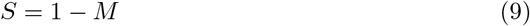

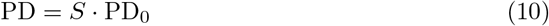

### 2.2 Parameter Calibration

A parameter set for Namdo (ND) cultivar was mostly based on existing parameter sets calibrated for Korean Mountain (KM) and Shantung Purple (SP). Only a few parameters were modified to reflect how ND grows in general compared to the other cultivars (Table 3). Minimum length of longest leaf (LM_min_) was increased to 100 cm which was the largest value found in our datasets. Potential maximum elongation rate (LER_max_) was accordingly adjusted to 5.56 cm d ^−1^ assuming full leaf expansion takes 18 days under optimal condition. Initial leaves at harvest (ILN) was set to 6 as found in a dissected seed garlic clove. Stay green (SG) was set to 1.5 d which was a value calibrated for SP cultivar. Storage temperature (ST) was assumed constant 8 °C during entire storage period. Seed bulb was assumed harvested on June 30th and storage duration (SD) was calculated accordingly. Note that it was our intention to select parameters that allow overestimation of biomass and leaf area for tracking potential growth while calibrating parameters to keep timing of phenology as close as possible to the observation.

**Table 3:** Parameters for leaf development

| Symbol | Value | Units | Description |
| --- | --- | --- | --- |
| $LTAR_{\max,a}$ | 0.4421 | $d^{-1}$ | Maximal leaf tip appearance rate asymptote |
| $LER_{\max}$ | 5.56 | $cm\ d^{-1}$ | Maximal leaf elongation rate |
| $LM_{\min}$ | 100 | cm | Minimum length of longest leaf |
| ILN | 6 | - | Initial leaf number |
| SG | 1.5 | - | Stay green |

We validated our new parameter set for ND cultivar using three datasets. The first dataset (D1) was collected from an experiment plot located at Research Institute of Climate Change and Agriculture (RICCA), Jeju, South Korea. ND cultivar was planted on October 8th, 2014 and harvested on June 19th, 2015. Growth and development measurements, such as leaf count, leaf area, biomass for each part, were recorded from mid-vegetative stage until harvest. This dataset was used for evaluating overall response of the ND parameter set we obtained above. The second dataset (D2) was collected from a temperature gradient greenhouse (TGG) located at the same site in Jeju. TGG is a glass house equipped with heaters on the one side of the wall that keeps a small, but constant gradient in the enclosed planting zones. Five planting zones were set up with 2 °C to 3 °C of temperature difference kept from end to end. ND cultivar was planted on October 7th, 2014 and harvested on May 17th, 2015. Similar measurements to the first dataset were recorded from early vegetative stage until harvest. This dataset was used for evaluating response of the model to modest temperature changes in a similar way that the model would be subject under future climate conditions. Scape was not removed in both datasets. The last third dataset (D3) was obtained from a farm field located at Jeongsil (JS) neighborhood in Jeju, South Korea. ND cultivar was planted on September 9th, 2009 and harvested on June 18th, 2010. Types of recorded variables were the same as other datasets. Scape was assumed to be removed shortly after its appearance out of the whorl. Only the visible portion of scape was cut and measured for biomass. Hourly time-series of weather data for each dataset were obtained from nearby Jeju station (184) operated by Korean Meteorological Administration.

### 2.3 Future Climate Projections

For assessing garlic yield changes in future climate conditions, we specifically chose two locations in South Korea (Figure 3). The first site was Gosan, Jeju located in a Southern island with a humid subtropical climate off the Korean peninsula, which is in Zone 9b in terms of USDA Hardiness Zones. Gosan is where Namdo (ND) cultivar is currently grown in a large scale for commercial purpose where stakeholders are interested to see how the cultivar would be performing and if any adjustments in growing practice such as if planting date shifts would be required in the long term. The second site was Chuncheon, Gangwon which is located in the middle of the Korean peninsula with a humid continental climate with cold winter. Chuncheon is in Zone 6b according to USDA Hardiness Zones and the current climate is not favorable for growing a Southern cultivar such as Namdo which has been historically more adapted to warm climate. However, it is not clear whether future climate conditions could allow growing of new crops in a region previously not suited for production. We assumed two RCP (Representative Concentration Pathway) scenarios, RCP4.5 and RCP8.5, for future climate projections (IPCC, 2014). Daily weather dataset for the two locations under RCP4.5 and RCP8.5 scenarios were obtained from AgClimate data portal (RDA, 2019) operated by Rural Development Administration in South Korea. Elevated CO_2_ concentrations for RCP scenarios were used accordingly for driving the integrated coupled gas-exchange model. For comparison with current climate conditions of each location, we also obtained 30-years normal weather data from 1980 to 2010 provided by the same data portal. The daily weather dataset was then scaled down to hourly time-series by applying an interpolation method specifically designed for different types of weather variables (Moon et al., 2019). Elevated CO_2_ levels for each year under future scenarios were obtained by linear interpolation between CO_2_ abundance projection reported in the Climate System Scenario Tables (IPCC, 2013).

**Figure 3:**
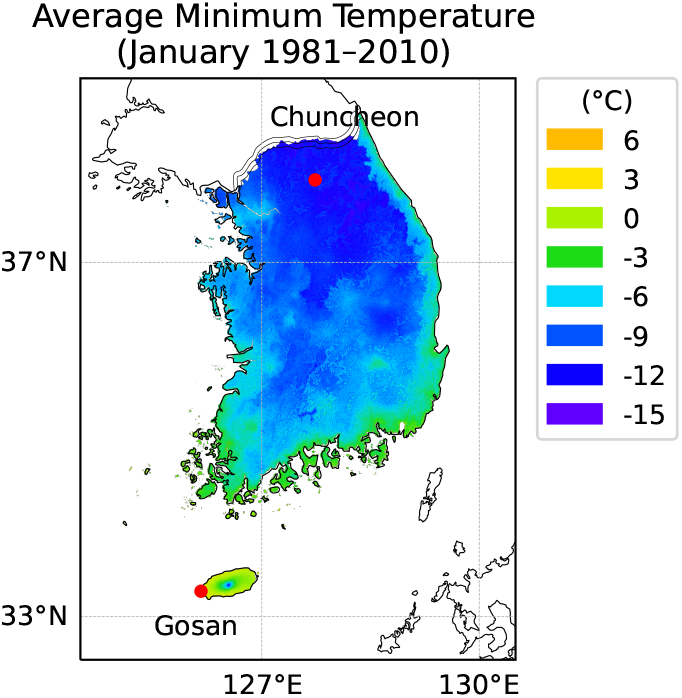
Geographic locations of Gosan and Jeju in South Korea. Gosan is located in a Southern island with a humid subtropical climate off the Korean peninsula. Chuncheon is located in the middle of the peninsula with a humid continental climate with cold winter. A color overlay represents average minimum temperature (°C) in January during past 30 years from 1981 to 2010 provided by National Institute of Agricultural Sciences, South Korea. According to USDA Hardiness Zones, Gosan is Zone 9b and Chuncheon is Zone 6b.

For each location, multiple runs of simulation were executed with combinations of treatments. In the case of future climate projections, 8 time windows from 2020s to 2090s with 10 years interval multiplied by 10 repetitions from different random seed produced 80 weather datasets for each RCP scenario. Historical weather was organized in a single time window referred to 1980s with 10 random repetitions, leading to 10 weather datasets. Planting date was adjusted from 240th to 350th DOY (day of the year) with 10 days interval, requiring 12 runs for each weather condition. In turn, the current normal scenario involved 120 runs of simulation while each RCP scenario required 960 runs. Immediate scape removal after appearance was assumed. Harvest date was set fixed to be May 15th considering practical harvest dates of ND cultivar grown in Jeju. Yield was based on fresh biomass of bulb at the harvest date. In model simulation, fresh biomass was calculated with 85 % moisture content in bulb. Optimal plating date was then determined by finding out a planting date which gives maximum yield for a given year. For the sake of yield comparison between planting dates, years were grouped into three time periods, namely 1980–2010s, 2020–2050s, and 2060–2090s. A data point in the 1980–2010 period was composed of 10 samples originated from a set of stochastic weather data pooled for the entire period whereas a data point in the future climate scenario was composed of 50 samples where the result of 10 stochastic weather datasets was combined with 5 periods of 10 years interval.

## 3 Results

### 3.1 Model Validation

#### 3.1.1 Dataset 1 (RICCA field)

Our parameter for ND cultivar was first evaluated with measurements from field grown garlic in dataset D1. Green leaf area from the model exhibited consistent overestimation with an average RMSE (root mean square error) of 275 cm^2^ while keeping a similar trajectory of leaf expansion and senescence pattern to the observation (Figure 4). The maximum green leaf area was estimated to be 1193 cm^2^ on April 28th while the maximum value in the dataset was 1150 cm^2^ observed on April 30th. The peak was reached in simulation shortly after scape appeared on April 23rd during early vegetative growth.

**Figure 4:**
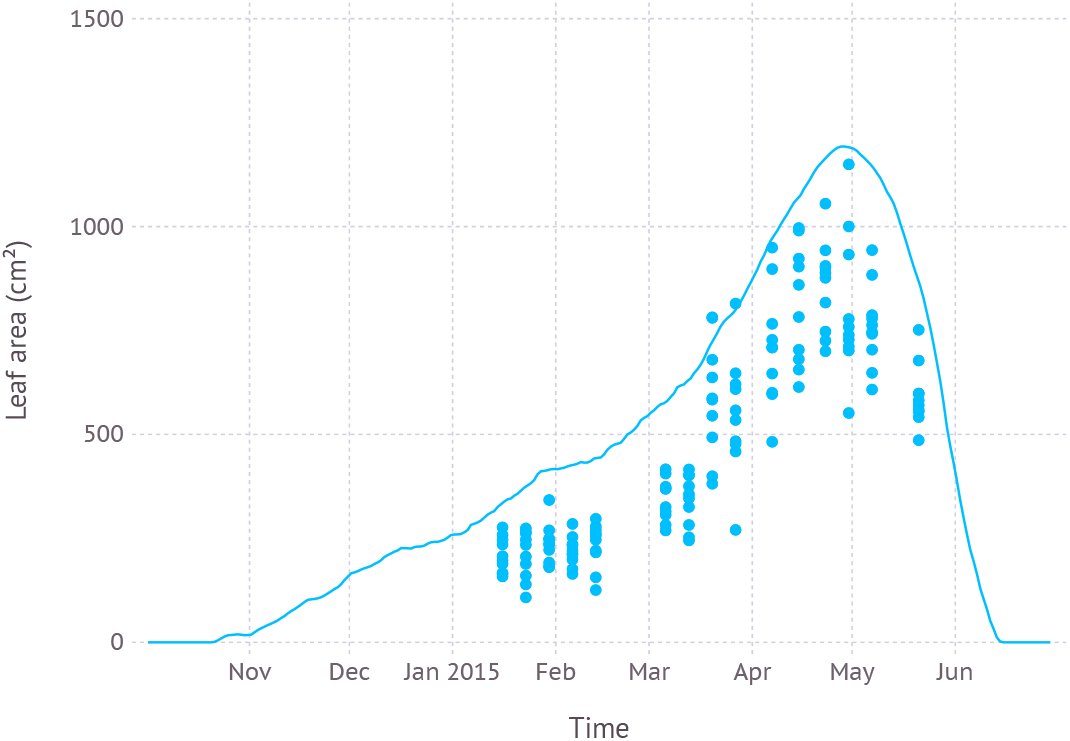
Comparison of model simulated green leaf area and observation from dataset D1 which was from field measurements and primarily used for evaluating new parameters.

Simulated biomass allocation to each part of plant structure, namely bulb, leaf, stalk, and scape, was close to the observation with an average RMSE of 5.0 g, 1.6 g, 2.5 g and 1.7 g, respectively (Figure 5). The maximum biomass of bulb from simulation was 22.6 g reached on June 14th while the observed maximum was 26.0 g measured on June 4th which was the final date recorded in the dataset. The maximum living leaf biomass from simulation was 8.4 g reached on April 27th and the maximum observed value was the same 9.9 g on April 30th. The maximum biomass of stalk, which consists of leaf sheath and scape, from simulation was 15.3 g close to observed maximum of 15.9 g. However, the timing of peak was shifted by almost three weeks later on May 27th compared to the observed May 7th. Since leaf sheath withered away at the end of growth stage, the only remaining part of living stalk would be a scape which was the reason why a convergence between two values occurred in mid-June. The simulated biomass of scape reached its maximum value of 9.5 g on June 20th which was almost identical to the observed maximum which, however, occurred earlier on June 4th. Note that the two slopes of biomass increase for simulation and observation were almost parallel to each other.

**Figure 5:**
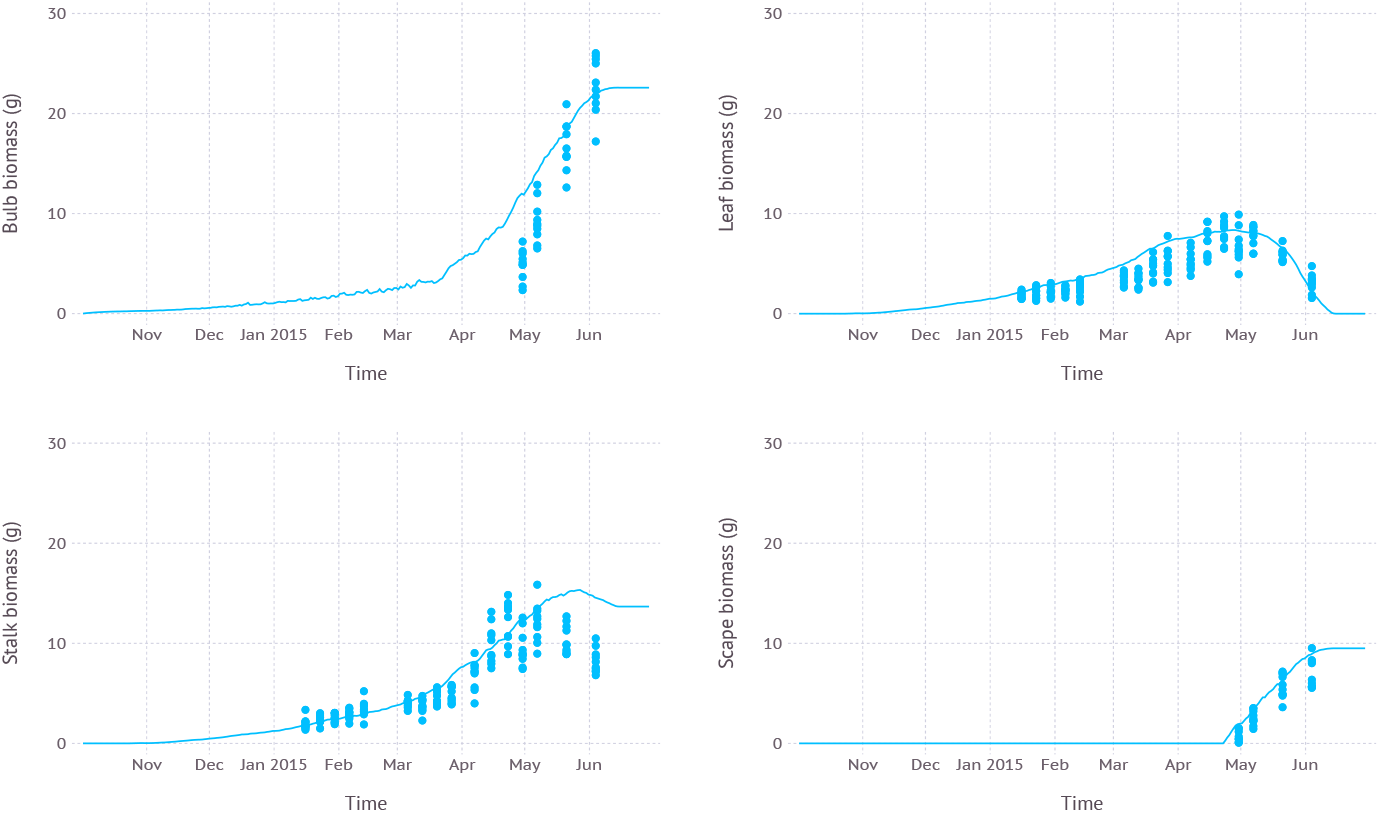
Biomass allocation to leaf, bulb, stalk, and visible portion of scape simulated over time compared with measurements from dataset D1 which was from field measurements and primarily used for evaluating new parameters.

Leaf development phenology simulated by the model was in agreement with fresh leaf count recorded in the dataset with an average RMSE of 1.6 leaves (Figure 6). According to the model, leaf initiation was completed on March 22nd with the total number of 16 leaves and the 16th leaf appeared on April 2nd. The final leaf was completely senesced and dropped after about 2.5 months of growth on June 15th.

**Figure 6:**
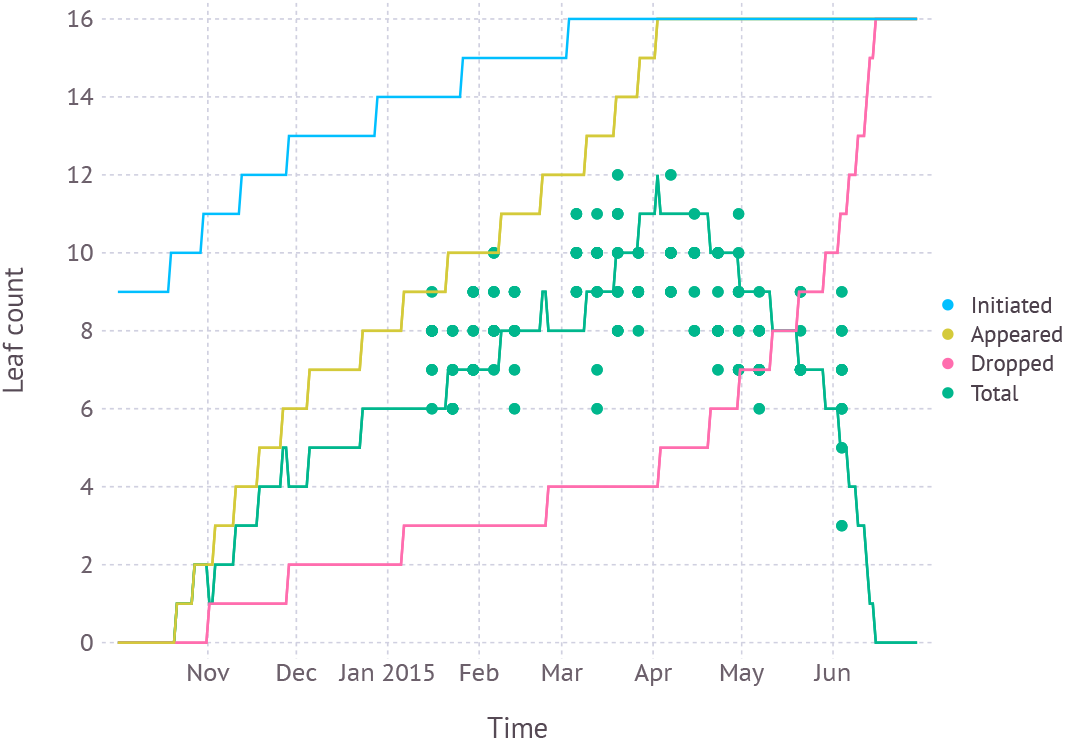
Simulated leaf development phenology compared with fresh leaf count recorded in dataset D1 which was from field measurements and primarily used for evaluating new parameters.

#### 3.1.2 Dataset 2 (RICCA TGG)

Plants grown in higher temperature zone 1 had larger green leaf area of 1521 cm^2^ and reached its peak earlier on April 20th when compared to plants grown in lower temperature zone 5 whose maximum green leaf area was estimated to be a smaller 1303 cm^2^ reached on May 9th (Figure 7). Simulation result was in agreement with observation recorded in dataset D2 in which zone 1 had larger leaf area of 1176 ± 239 cm^2^ on April 17th, in contrast to zone 5 having smaller leaf area of 1124 ± 90 cm^2^ on the same date. After three weeks on May 9th, the difference was flipped over where plants in zone 1 had senesced faster, resulting into a smaller green leaf area of 512 ± 55 cm^2^ while zone 5 was still greener with 701 ± 70 cm^2^. Mean temperature recorded in dataset D2 for zone 1 was 15.5°C and zone 5 was 12.6 °C.

**Figure 7:**
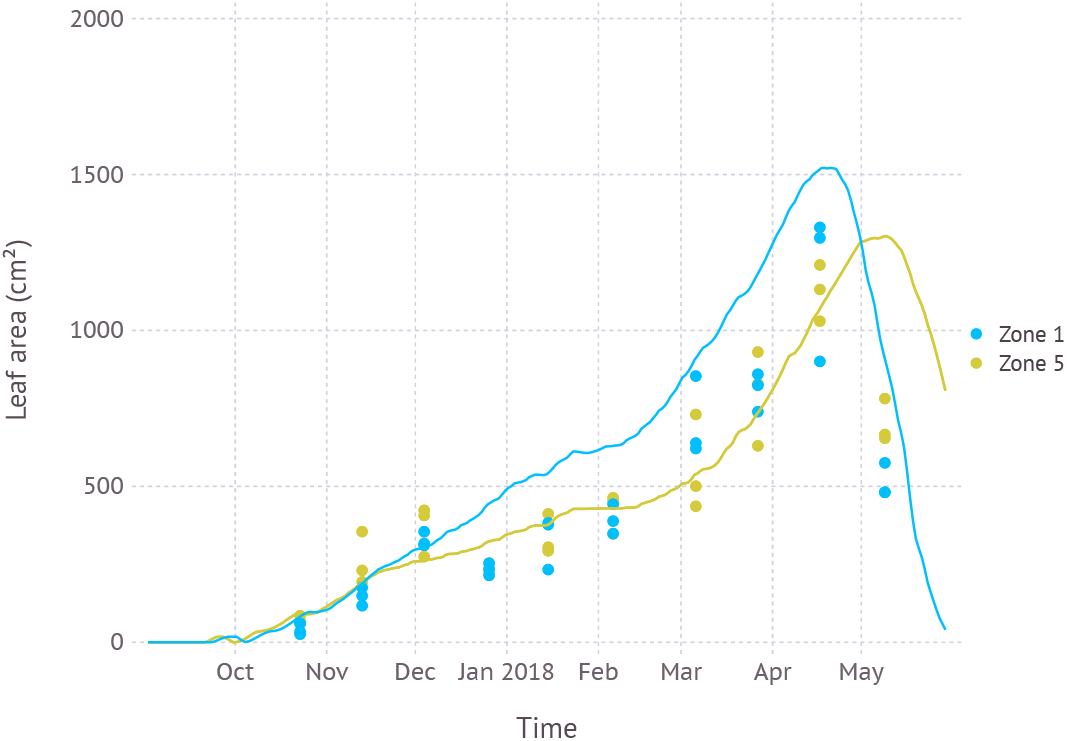
Comparison of model simulated green leaf area and observation for two temperature zones (Zone 1: high, Zone 5: low) recorded in dataset D2 from a temperature-controlled glass house.

#### 3.1.3 Dataset 3 (JS field)

Simulated biomass allocation compared with another independent dataset D3 came with an average RMSE of 7.4 g, 3.6 g and 4.2 g for bulb, leaf, and stalk, respectively (Figure 8). The maximum bulb biomass from simulation was 37.4 g reached on June 22nd while the observed maximum was 28.7 g measured on May 26th. The maximum living leaf biomass from simulation was 10.0 g reached on May 1st and the maximum observed value was 9.1 g on April 28th. The maximum biomass of stalk from simulation was 13.5 g on May 1st when scape just appeared and subsequently removed. The maximum stalk biomass recorded in the dataset was 12.3 g observed on April 28th. Once the scape was removed and no longer a part of stalk composition, stalk biomass gradually decreased as the mature sheath stopped growing and started senescence. According to the dataset, scape was removed during reproductive stage, but an exact date of the removal was not recorded. Thus we assumed scape removal took place as soon as a tip of the scape became visible out of the whorl which occurred on May 1st by model estimation. The scape biomass at the time of removal was 5.7 g.

**Figure 8:**
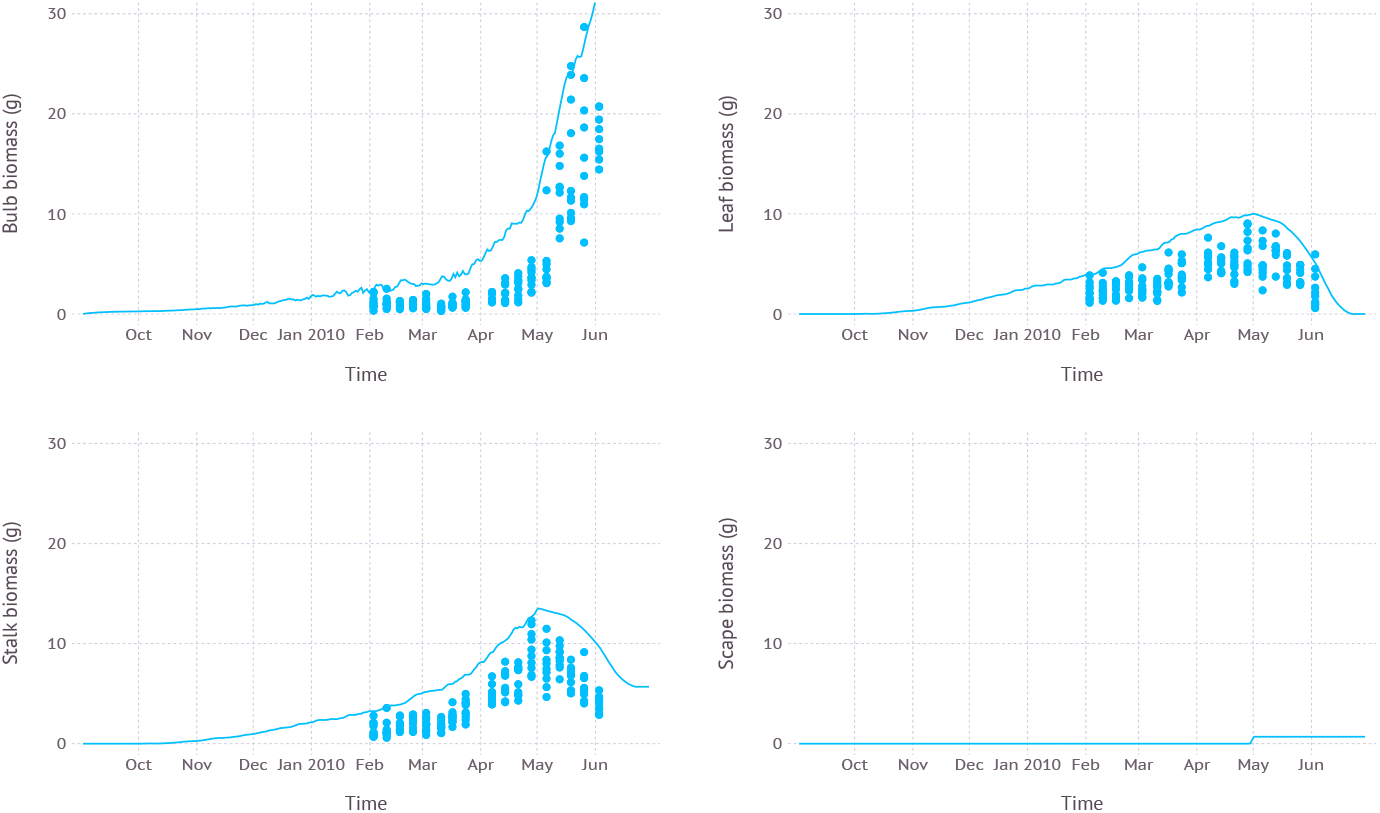
Biomass allocation to leaf, bulb, stalk, and visible portion of scape simulated over time compared with measurements from dataset D3 which was from field measurements with scape removal.

### 3.2 Yield Projection

#### 3.2.1 Gosan

In the current climate condition, fresh yield on ND cultivar in Gosan, Jeju was estimated to be maximum at an average of 6.8 kg m^−2^ with standard deviation of 0.8 kg m^−2^ when planted in late August (240 DOY) which was closely followed by early September planting (260 DOY) with an average of 6.7 ± 0.7 kg m^−2^ (Figure 9). A similar level of high yield was maintained until late September then yield gradually decreased with later planting dates. The difference between maximum yield from early planting date and minimum yield from later planting date was 3.7 kg m^−2^. In the near future from 2020s to 2050s, a similar pattern was observed that high yield was achieved with early planting in September. An overall yield was increased to 7.5±0.8kg m^−2^ under RCP4.5 and 7.6±0.8kg m^−2^ under RCP8.5 scenario when compared to the current climate. There was no clear difference between RCP4.5 and RCP8.5 scenarios in terms of estimated yield for a given planting date. In the distant future from 2060s to 2090s, RCP4.5 scenario still maintained a similar pattern where high yield was achieved in early planting dates, but then the range of potential high yield was expanded to later planting dates. For example, planting dates from late August (240 DOY) to mid-October (290 DOY) all resulted in an average yield closely ranged from 7.6 kg m^−2^ to 7.7 kg m^−2^. The difference between maximum and minimum yield estimated in the range of planting dates was reduced to 3.0kg m^−2^. Under RCP8.5 scenario, the yield curve was more flattened that the difference was only 1.8 kg m^−2^ between all planting dates. The maximum yield was 8.3 ± 0.6 kg m^−2^ in early-October (280 DOY).

**Figure 9:**
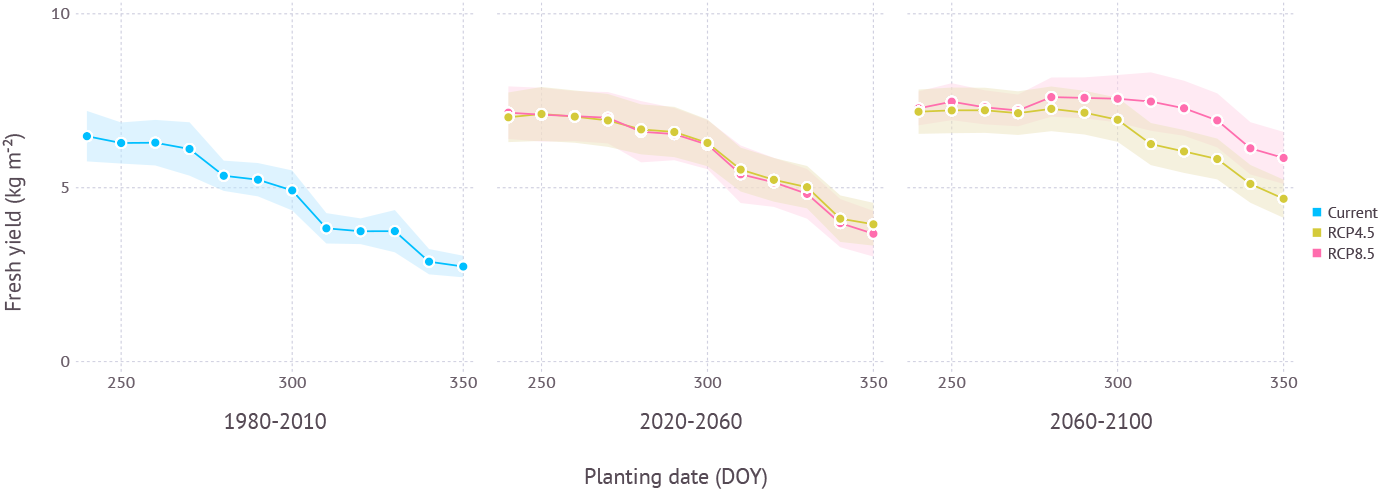
Yield estimation within a range of planting dates under current and future climate projection in Gosan, Jeju which is a region grows ND cultivar at commercial scale. The current climate indicates simulation results with 30-years normal weather data from 1980–2010. Shades represent a range of ±1 standard deviation from mean fresh yield.

When optimal plating dates were assessed from current to future climate conditions by 10 year intervals, a clear trend of shifting towards later planting date was found in the future projection although its strength varied depending on scenarios (Figure 11). Both scenarios began with optimal planting date in early to mid-September then showed a strong divergence at the end of the century that the optimal planting date surfaced from late September to early October under RCP4.5 scenario, while mid-November became a possibility under RCP8.5 scenario.

#### 3.2.2 Chuncheon

In the current climate condition, yield of ND cultivar in Chuncheon, Gangwon was estimated close to nil with a maximum yield 0.06 ± 0.05 kg m^−2^ due to impeded leaf growth and increased mortality from cold winter temperature during early vegetative growth (Figure 10). However, under future projections, yield became more viable thanks to warmer climate. In the near future from 2020s to 2050s, maximum estimated yield was 1.5±0.7 kgm^−2^ under RCP4.5 and 1.3 ± 0.9 kg m^−2^ under RCP8.5 scenario when both planted in early September (250 DOY). In the distant future from 2060s to 2090s, more yield was achievable with 2.5±0.8 kg m^−2^ under RCP4.5 and 4.3± 1.1 kgm^−2^ under RCP8.5 scenario with early to mid-September plating dates, 250 DOY and 240 DOY, respectively. The difference between maximum and minimum yield estimated in the range of planting dates was 0.6 kg m^−2^ in early period and 1.0 kg m^−2^ in late period under RCP4.5 scenario. RCP8.5 scenario initially had a similar range of estimated yield over multiple planting dates with 0.5 kg m^−2^ in early period, but then showed much higher variance of 1.7 kg m^−2^ in late period.

**Figure 10:**
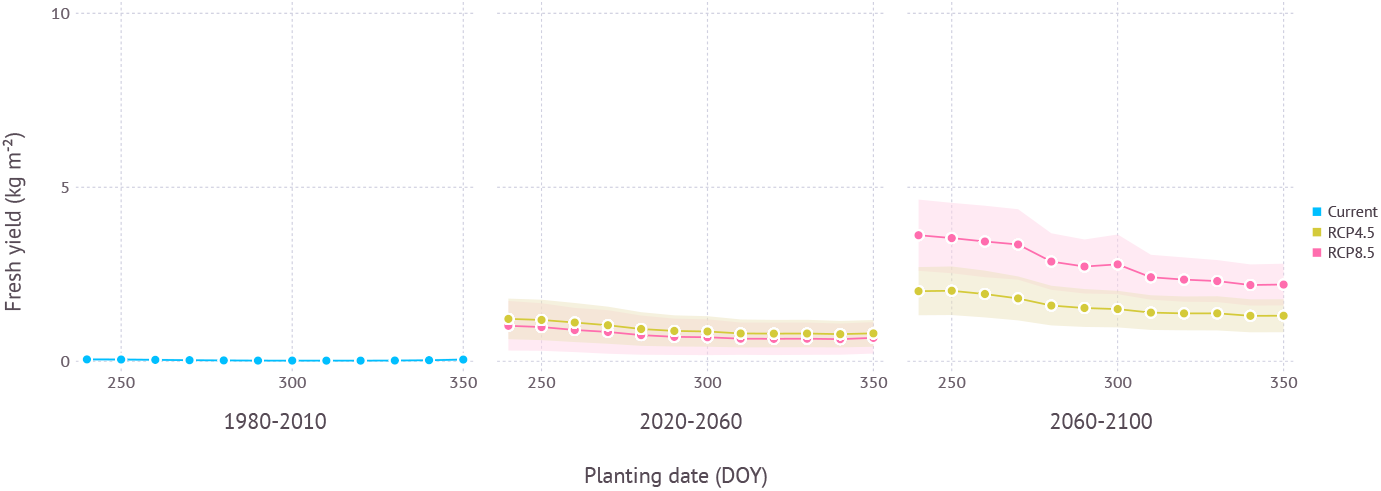
Yield estimation within a range of planting dates under current and future climate projection in Chuncheon, Gangwon where current climate is not favorable for growing ND cultivar. The current climate indicates simulation results with 30-years normal weather data from 1980–2010. Shades represent a range of ±1 standard deviation from mean fresh yield.

In terms of optimal planting dates for each period, planting in around early September consistently turned out to work best regardless of climate scenarios and estimated yield gradually declined with later planting dates in all scenarios (Figure 11).

**Figure 11:**
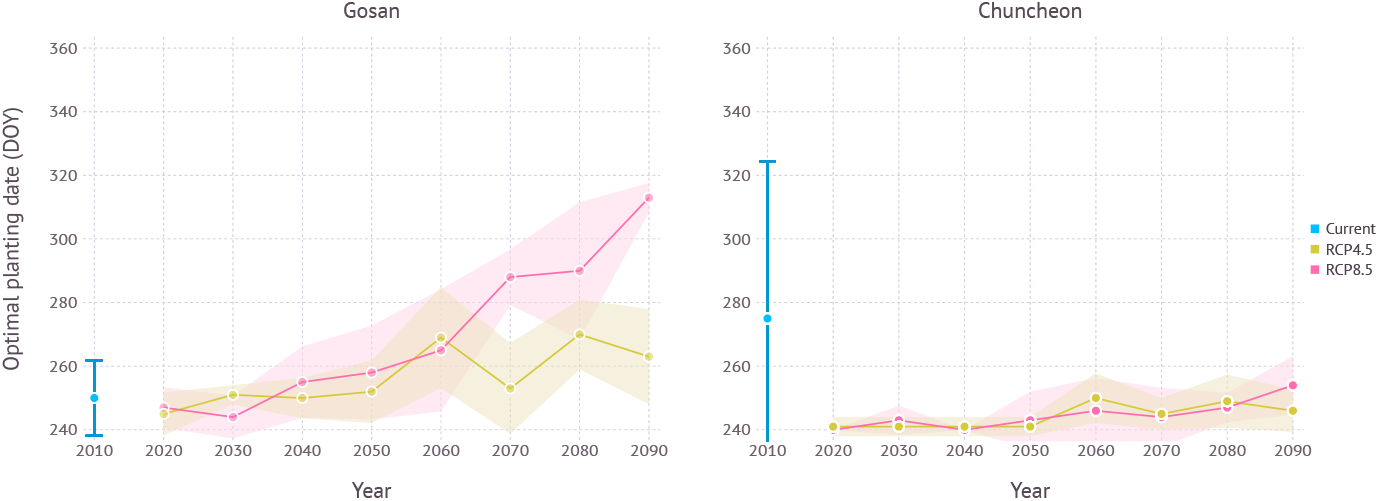
Optimal planting dates estimated for each year under current and future climate projections in Gosan and Chuncheon. Year 2010 indicates simulation results with 30-years normal weather data from 1980–2010 representing current climate. For RCP scenarios, each year indicates 10-year average, *e.g.*, 2090 refers to 2090–2100. Error bar and shades represent a range of ±1 standard deviation from mean optimal planting date.

## 4 Discussion

### 4.1 Model Validation

#### 4.1.1 Parameter Calibration

Our parameter set for ND cultivar was mostly derived from existing parameters for Korean Mountain (KM) and Shantung Purple (SP) with small changes (Hsiao et al., 2019). The final parameter set used in our simulation turned out to be similar to one calibrated for SP cultivar. It would not be unexpected given that ND was originated from a Chinese cultivar and hence there is a chance SP might be also related with this original cultivar due to geographical proximity (Kim et al., 2009).

#### 4.1.2 Overestimation

Simulation with the calibrated parameter set often resulted in overestimation in a sense that model output tends to touch upper boundaries of observed data points (Figure 4,5,7,8). This overestimation is intended and expected because our model estimates for biomass and yield potential for a given environmental condition with assuming no limiting factors such as water and nutrient while the field conditions are likely less than optimal. Some types of overestimation also presumably came from difference between physical measuring methods and modeled algorithms. For example, green leaf area recorded in the dataset was measured by taking only non-withered leaves at the time of observation which is prone to lose leaves with some green parts intact. On the other hand, green leaf area from the model was calculated for individual leaves at a fractional scale by tracking current green portion of the leaf for each time step. These difference could have been one of the reasons leading to overestimation in leaf area simulation (Figure 4,7).

#### 4.1.3 Temperature Regimes

Evaluation of model response under slightly separate temperature regimes as conducted in temperature gradient house provided an insight how elevated air temperature under RCP scenarios would affect plant growth (Figure 7). In short, plant will grow larger and faster and die earlier in warmer conditions. An average of 3 ° C difference in temperature led to more than two weeks of shift in growth peak and 15% change in total green leaf area. Subsequent changes in senescence timing would imply a need for finding out optimal harvest dates which we assumed as a rather constant management decision in our simulation.

#### 4.1.4 Storage Condition

Storage duration (SD) affects maximum leaf tip appearance rate (LTARmax) via the dynamic phyllochron model (Equation 1,2,3). Storage temperature (ST) along with storage duration then decides number of leaves initiated inside a seed bulb at the time of planting. Despite their importance on phenology, there often exits no record on when seed bulbs were harvested and in which condition they were stored until the date of planting. With controlled experiments on storage condition, we could have better understanding on how leaves are initiated during storage period and whether more sophisticated approaches like a dynamic plastochron would be necessary.

#### 4.1.5 Other Environmental Cues

While our assessment for yield projection under future condition was primarily driven by model response to temperature regimes, there is still room for considering other environmental cues. For instance, we did not have a separate vernalization process in the model, but assumed winter chilling was always at an adequate level for triggering continuous development in the following warm spring. While it worked reasonable well in most cases, some processes like mortality from cold stress could capitalize on this additional cue to make dormant plants less vulnerable to cold damage, for instance. Soil water availability would be another important cue in terms of assessing irrigation requirement under future climate conditions given that garlic plots are often irrigated to prevent water deficiency during reproductive stage.

### 4.2 Yield Projection

#### 4.2.1 Gosan

Jeju, including Gosan, is a region where southern-type garlic cultivars like Namdo (ND) has been grown at commercial scale thanks to warm climate. Northern regions in Korean peninsula often have colder winter which prevents growing southern-type cultivars. Growers instead choose northern-type cultivars for cold hardiness. As southern cultivars have advantage of higher yield and shorter growing season, whether they could be adapted to northern regions has been an important question to many growers and stakeholders. According to our simulation results using a process-based model, ND cultivar would continue growing well with a slightly higher yield (Figure 9). The optimal planting date in early September estimated for the current climate condition was indeed close to the current practice held in Jeju. This practice is, however, expected to be delayed in the near future if growers keep trying to maximize yield under both climate scenarios (Figure 11). The amount of planting date delay depends on the scenarios and the model estimated that RCP4.5 scenario could see a shift less than a month while RCP8.5 scenario could potentially result in two months of late planting. Note that although optimal planting dates were seemingly changed drastically, yield estimation curve versus planting date was actually getting more flattened and the difference between maximum and minimum yield became smaller in warmer climate conditions. Such changes imply that growers would have more options to choose planting date better suits their own needs. For example, October planting in the future would still promise a yield level close to potential maximum while taking a relatively shorter growing season to help reducing overall cost of labor and resources. High yield does not necessarily lead to high profitability when longer growing season increases overall water demand entailed by higher irrigation cost for compensation (Lobell, 2014). Looking further in distant future, it was more clear to expect a shift in optimal planting to later dates although yield difference to earlier planting dates was not significant.

#### 4.2.2 Chuncheon

Chuncheon is located in the middle of Korean peninsula and its continental climate frequently experiences below freezing temperature during winter season. Hence only northern-type cultivar adapted to cold climate have been grown in this region. Our simulation results confirmed expected yield for the current climate was almost non-existent as most plants subject to mortality (M) could not survive after episodes of extreme cold. By contrast, future climate conditions were projected to become more favorable in both scenarios so that expecting some tangible yields would be at least feasible (Figure 10). Generally early planting dates were favored in terms of optimal yield as similar in the current growing region like Jeju (Figure 11). Although the level of estimated yield was still lower when compared to the yield currently obtainable in other established regions, growers may be able to take advantage of this new opportunity for expanding crop portfolio during winter season.

### 4.3 Climate Adaptation

It is important to tease apart an effect of adaptation introduced by planting date shift in the future from an impact of planting date shift in the current condition (Lobell, 2014; Challinor et al., 2018). According to the result from Gosan, planting date shift in established growing region has little effect on adaptation as evidenced by a similar or smaller range between maximum and minimum yield when comparing current and future climate scenarios (Figure 9). Higher estimated yield was a result of increased productivity under more favorable condition primarily due to higher temperature and elevated CO_2_, but not from potential phenological changes occurred by planting date shift. In non-established regions like Chuncheon, however, an effect of adaptation increased virtually from zero to a substantial amount when transitioning to new climate condition as evidenced by a rising slope of yield estimation curve (Figure 10). In other words, the importance of planting date as a climate adaptation strategy depends on how crops are adapted to the local condition.

Underlying uncertainties of crop models often hinder model driven assessment of climate change adaptation (Rosenzweig et al., 2014; Holzkämper et al., 2015; Corbeels et al., 2018). We adopted a process-based model with coupled gas-exchange to minimize uncertainties in physiological responses to temperature and CO_2_, but there still remain many processes that can use improvements. For instance, water stress response was solely dependent on leaf water potential disconnected with soil water status albeit a less practical implication when garlic plants are generally irrigated to meet high water demand during reproductive stage. Nitrogen response is another important factor that the model currently does not take into account and instead assume non-limiting due to extensive use of fertilizers in practice, but will be critical for assessing an exact cost of production in finding optimal yield.

## Conflict of Interest Statement

The authors declare that the research was conducted in the absence of any commercial or financial relationships that could be construed as a potential conflict of interest.

## Funding

The information, data, or work presented herein was funded in part by the Cooperative Research Program for Agricultural Science and Technology Development, Rural Development Administration, Republic of Korea under Grant Number PJ015124012021. The views and opinions of authors expressed herein do not necessarily state or reflect those of the funding agency.

## Notes

https://github.com/cropbox/Garlic.jl

### Competing Interest Statement

The authors have declared no competing interest.

